# *MEOX2* homeobox gene promotes growth of malignant gliomas

**DOI:** 10.1101/2021.08.24.457477

**Authors:** Anna Schönrock, Elisa Heinzelmann, Bianca Steffl, Ashwin Narayanan, Damir Krunic, Marion Bähr, Jong-Whi Park, Claudia Schmidt, Koray Özduman, M. Necmettin Pamir, Wolfgang Wick, Felix Bestvater, Dieter Weichenhan, Christoph Plass, Julian Taranda, Moritz Mall, Şevin Turcan

## Abstract

Glioblastoma (GBM) is an aggressive tumor that frequently exhibits gain of chromosome 7, loss of chromosome 10 and aberrantly activated receptor tyrosine kinase signaling pathways. Here, we identify mesenchyme homeobox 2 (MEOX2) on chromosome 7 with increased expression in GBM as a salient oncogenic transcription factor. Specifically, we show that MEOX2 overexpression leads to increased ERK phosphorylation, and we identify a phosphorylation site on MEOX2 that regulates its transcriptional activity by altering its subnuclear localization. We show that MEOX2 overexpression can lead to increased growth in GBM implantation models and cooperates with loss of p53 and PTEN in cerebral organoid models of human malignant gliomas to induce cell proliferation. Furthermore, using high-throughput genomics, we identify transcriptional target genes of MEOX2 in patient-derived GBM tumorsphere models and a fresh frozen GBM tumor. These analyses show that MEOX2 activates several oncogenic pathways involved in MAPK signaling and extracellular matrix organization. Furthermore, MEOX2 binds to oncogenic ETS factors and known glioma oncogenes such as *FABP7*. In total, we reveal a novel role for MEOX2 in GBM initiation and progression and demonstrate that MEOX2 can enhance ERK signaling through a feed-forward mechanism.

**Significance Statement:** Glioblastoma (GBM) harbors gain of chromosome 7 as an early driver event. In this study, we show that mesenchyme homeobox 2 (MEOX2), an aberrantly upregulated transcription factor on chromosome 7, is an oncogene in human glioblastoma. In contrast to GBM, *MEOX2* expression is very low in normal brain. We show that MEOX2 cooperates with p53 and PTEN loss to promote tumor initiation in cerebral organoid models. In addition, we identify direct and indirect molecular targets of MEOX2 and demonstrate its role in activating the ERK signaling cascade. These findings identify a novel oncogene in GBM and highlight the transcriptional networks hijacked by these tumors to activate signaling pathways central to GBM biology.

## Introduction

Glioblastoma multiforme (GBM) is a malignant tumor of the central nervous system with a high proliferative capacity and a diffuse pattern of brain invasion. Despite aggressive multimodal treatment, only about 3-5% of patients survive three years or longer. Several core signaling pathways are activated in GBM, and include the p53, the receptor tyrosine kinase (RTK)/Ras/phosphoinositide 3-kinase (PI3K), and retinoblastoma (Rb) signaling pathways (Cancer Genome Atlas Research 2008). Common genetic alterations in GBM include gain of chromosome 7, loss of chromosome 10, alterations in *TP53*, epidermal growth factor receptor (*EGFR*) and platelet-derived growth factor receptor (*PDGFR*), and mutations in mouse double minute homolog 2 (*MDM2*) and the phosphatase and tensin homolog (*PTEN*) gene (Brennan et al. 2013). These molecular alterations act together to promote GBM growth, survival, escape from cell-cycle checkpoints, senescence, and apoptosis.

Aberrantly overexpressed transcription factors have the potential to drive GBM (Suva et al. 2014; Singh et al. 2017). These observations raise the possibility that overexpression of key transcription factors may be sufficient to drive a malignant GBM phenotype or that they interact with early driver molecular alterations to initiate tumors. Previously, we identified *MEOX2* as a transcription factor highly expressed in IDH wild-type gliomas and a recent study has associated *MEOX2* expression with poor GBM prognosis (Turcan et al. 2018; Tachon et al. 2019). *MEOX2* is localized to chromosome 7, and recurrent gain of chromosome 7 and loss of chromosome 10 are common GBM features that occur early in gliomagenesis (Ozawa et al. 2014). Several genes on chromosome 7, such as platelet-derived growth factor (*PDGFA*) and homeobox A5 (*HOXA5*), have been shown to drive tumor aggressiveness and radio-resistance (Ozawa et al. 2014; Cimino et al. 2018).

MEOX2 is a homeobox-containing transcription factor that plays an essential role in developing tissues (LePage et al. 1994; Mankoo et al. 1999). It activates p16 and p21 through DNA-dependent and independent mechanisms, and increased MEOX2 expression leads to cell cycle arrest and endothelial cell senescence (Smith et al. 1997; Douville et al. 2011). In cancer, the role of MEOX2 is poorly defined and likely context-dependent, acting either as a tumor suppressor or as an oncogene (Ohshima et al. 2009; Cao et al. 2010).

Here, we hypothesized that MEOX2 could provide a selective advantage in GBM. We demonstrate that MEOX2 overexpression increases tumor growth *in vivo* and leads to a significant proliferative advantage in human cerebral organoid models of GBM, highlighting the potential role of MEOX2 as an early driver of gliomagenesis. Mechanistically, we find a feedforward loop in which gene regulatory activity of MEOX2 is likely controlled by its phosphorylation level, which in turn leads to increased ERK signaling. Our results suggest that MEOX2 is one of key oncogenes on chromosome 7 that is co-opted by GBM to drive gliomagenesis.

## Results and Discussion

### MEOX2 is highly expressed in IDH wild-type GBM

Analysis of the combined lower-grade and GBM molecular profiling datasets from The Cancer Genome Atlas (TCGA) revealed a significantly higher *MEOX2* expression in IDH wildtype GBM (Fig. 1A; Supplemental Fig. 1A-C). Because a role for MEOX2 in endothelial cells has been defined previously (Wu et al. 2005), we interrogated published single-cell RNA-sequencing (RNA-seq) datasets in adult GBMs to determine the cell populations with higher MEOX2 levels. Analysis of the data by Neftel and colleagues. (Neftel et al. 2019) showed that *MEOX2* expression in GBM was restricted to the tumor cells (Fig. 1B). In comparison, *MEOX2* expression in the normal brain is undetectable or very low (Supplemental Fig. 1D).

**Figure 1.**
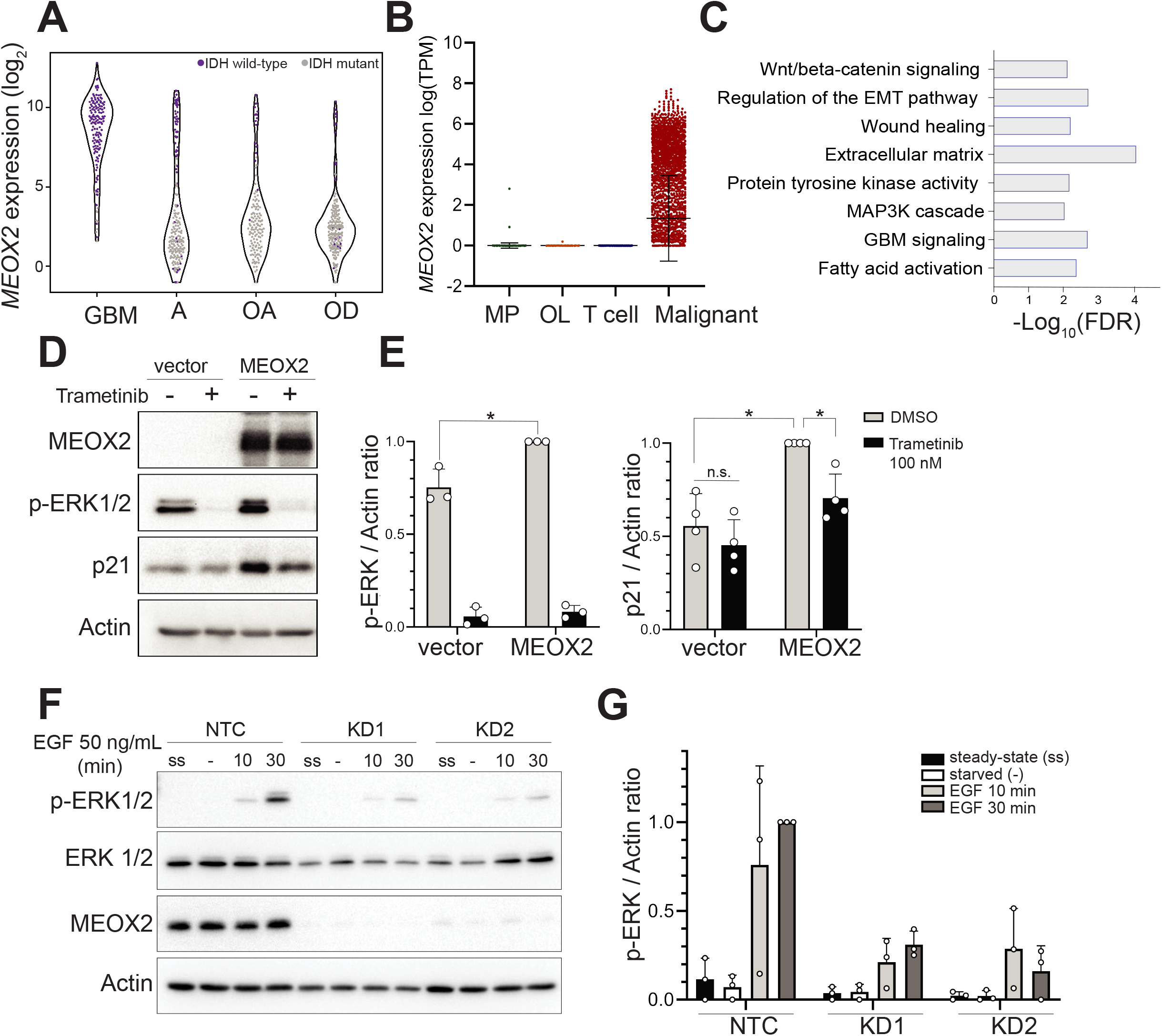
*MEOX2* is expressed in GBM and induces phosphorylation of ERK. (A) Analysis of MEOX2 gene expression obtained from The Cancer Genome Atlas (TCGA) database of lower grade gliomas and GBM including IDH mutant (gray) and IDH wild-type (purple) tumors. Data downloaded from GlioViz (Bowman et al. 2017). GBM, glioblastoma; A, astrocytoma; OA, oligoastrocytoma; OD, oligodendroglioma. (B) MEOX2 expression in published GBM single-cell RNA-seq data (Neftel et al. 2019) assigned to macrophages (*n* = 536), oligodendrocytes (*n* = 210), T cells (*n* = 80) and malignant tumor cells (*n* = 4916). Mean ± SD; two-sided Welsh’s test. (C) Pathway enrichment analysis of the top 50 genes correlated with MEOX2 expression in the TCGA dataset. (D) Western blot analysis of MEOX2, phospho-ERK (p-ERK1/2), and p21 in HEK293TN cells transiently transfected with empty vector or MEOX2 for 48 hours. Transfected cells were treated with 100 nM trametinib (+) or control DMSO (-) for 24 hours. (E) Quantification of p-ERK relative to total actin levels (left), quantification of p21 relative to actin levels (right). Mean ± SD, *n* = 3 (left), *n* = 4 (right), two-sided Welsh’s test. (F) p-ERK levels in TS667 GBM tumorsphere line transduced with non-targeting control (NTC) or one of two single guide RNAs targeting MEOX2 (KD1, KD2). Cells were either kept under steady-state (ss) condition or starved (-) for 48 hours and stimulated with 50 ng / mL EGF for 10 or 30 min (10, 30). (G) Quantification of (F) showing p-ERK relative to actin in NTC, KD1 and KD2 TS667 cell lines. Mean + SD. *n* = 3. Two-sided Welsh’s test).

The transcriptional effects of MEOX2 in GBM are unknown, we therefore aimed to elucidate the pathways that are affected by aberrant MEOX2 activity in GBMs. We identified the top 50 genes that positively correlated with *MEOX2* expression in the TCGA GBM dataset (Supplemental Table 1). Gene programs involving the ERK1/2 cascade (e.g., *SPRY2, EGFR, PDGFA, P2RY1, PLA2G5, SPRY1, FGFR3, SPRY4*) and fatty acid synthesis (*ELOVL2, ACSL3, PLA2G5*) significantly correlated with *MEOX2* expression (Supplemental Table 2). To determine the pathways associated with MEOX2 in GBMs, we stratified the TCGA IDH-wildtype GBM tumors into MEOX2 high and low cohorts and identified the significantly deregulated pathways (Supplemental Table 3). Among the upregulated genes, pathways including protein tyrosine kinase (RTK) activity, extracellular matrix, GBM signaling, and regulation of the epithelial-mesenchymal transition (EMT) pathway were significantly enriched (Fig. 1C). Based on these findings, we hypothesized that MEOX2 could act as an oncogene in GBM to, in part, amplify the signaling output by RTKs.

### MEOX2 induces phosphorylation of ERK *in vitro*

MEOX2 is a homeobox transcription factor with little known about its function and regulation. Given that MEOX2 expression correlates with oncogenes such as EGFR and hallmark oncogenic pathways such as RTK signaling and EMT in GBM (Fig. 1C), we wondered whether MEOX2 may co-operate with signaling pathways activated by RTKs. To assess its function, we overexpressed MEOX2 in 293 or 293TN cells. Overexpression of MEOX2 resulted in increased p21 protein levels, a known MEOX2 downstream target, along with a significant increase in ERK phosphorylation (p-ERK) (Fig. 1D, E). Treatment of 293TN cells with the MEK inhibitor trametinib led to a decrease in p21 and p-ERK levels (Fig. 1D, E). This finding suggests that the transcriptional activity of MEOX2 could be regulated by its phosphorylation status. We next sought to determine whether MEOX2 impacts ERK phosphorylation in GBM models. CRISPR/Cas9-mediated knockdown of MEOX2 using two single-guide RNAs (KD1, KD2) in two patient-derived GBM tumorsphere lines, TS600 and TS667, showed significant reduction in p-ERK in the TS667 line upon EGF stimulation, whereas p-ERK levels in the TS600 line were not altered by reduced MEOX2 expression (Fig, 1F, G; Supplemental Fig. 2A, B). These results suggest that MEOX2 can regulate ERK phosphorylation in GBM models. However, regulation may depend on baseline ERK activity since in the EGFR-amplified TS600 line with higher steady state p-ERK, MEOX2 did not affect p-ERK levels.

### MEOX2 transcriptional activity is regulated by phosphorylation

Our finding that MEK inhibition reduced MEOX2-mediated induction of p21 (Fig. 1D) suggests that ERK might phosphorylate MEOX2. Indeed, we identified ERK-dependent phosphorylation sites upstream of the homeobox-domain of MEOX2 in patient-derived GBM tumorspheres by mass spectrometry following immunoprecipitation of endogenous or overexpressed MEOX2 (Fig. 2A). Particularly, the Ser^155^ harbors a consensus MAPK motif (S/T)P and has been reported in a large-scale phospho-proteomics analysis of breast cancer (Mertins et al. 2016), which led us to consider whether this may be a regulatory phosphorylation site in MEOX2. To determine whether Ser^155^ could lead to altered activity of MEOX2, the serine residue was changed to alanine (S155A). MEOX2^S155A^ led to decreased ERK1/2 phosphorylation compared to wild-type MEOX2 upon overexpression, yet both increased ERK1/2 phosphorylation compared to baseline levels (Fig. 2B, C). Importantly, p21, the transcriptional target of MEOX2, was only slightly induced in MEOX2^S155A^ overexpressing cells, such that p21 levels were similar to trametinib treated MEOX2-overexpressing cells. (Fig. 2B, D). S155A substitution had no effect on protein stability (Fig. 2E, F), so we investigated whether the nuclear distribution of the phospho-mutant protein may be altered. Using maximum intensity projection of *z* stack images, we analyzed the distribution of fluorescence signal around the nuclear envelope compared to the whole nucleus (Fig. 2H). Indeed, MEOX2^S155A^ showed altered nuclear localization with a significantly increased distribution in the nuclear envelope and nuclear lamina area compared with the whole nucleus (Fig. 2H, I). Non-activated transcription factors have been shown to be sequestered in the inner nuclear envelope (Heessen and Fornerod 2007). Together with the observation that MEOX2^S155A^ triggered lower p21 expression compared with the wild-type protein (Fig. 2D, G), these results suggest that phosphorylation could regulate the nuclear sublocalization and in turn the transcriptional activity of MEOX2.

**Figure 2.**
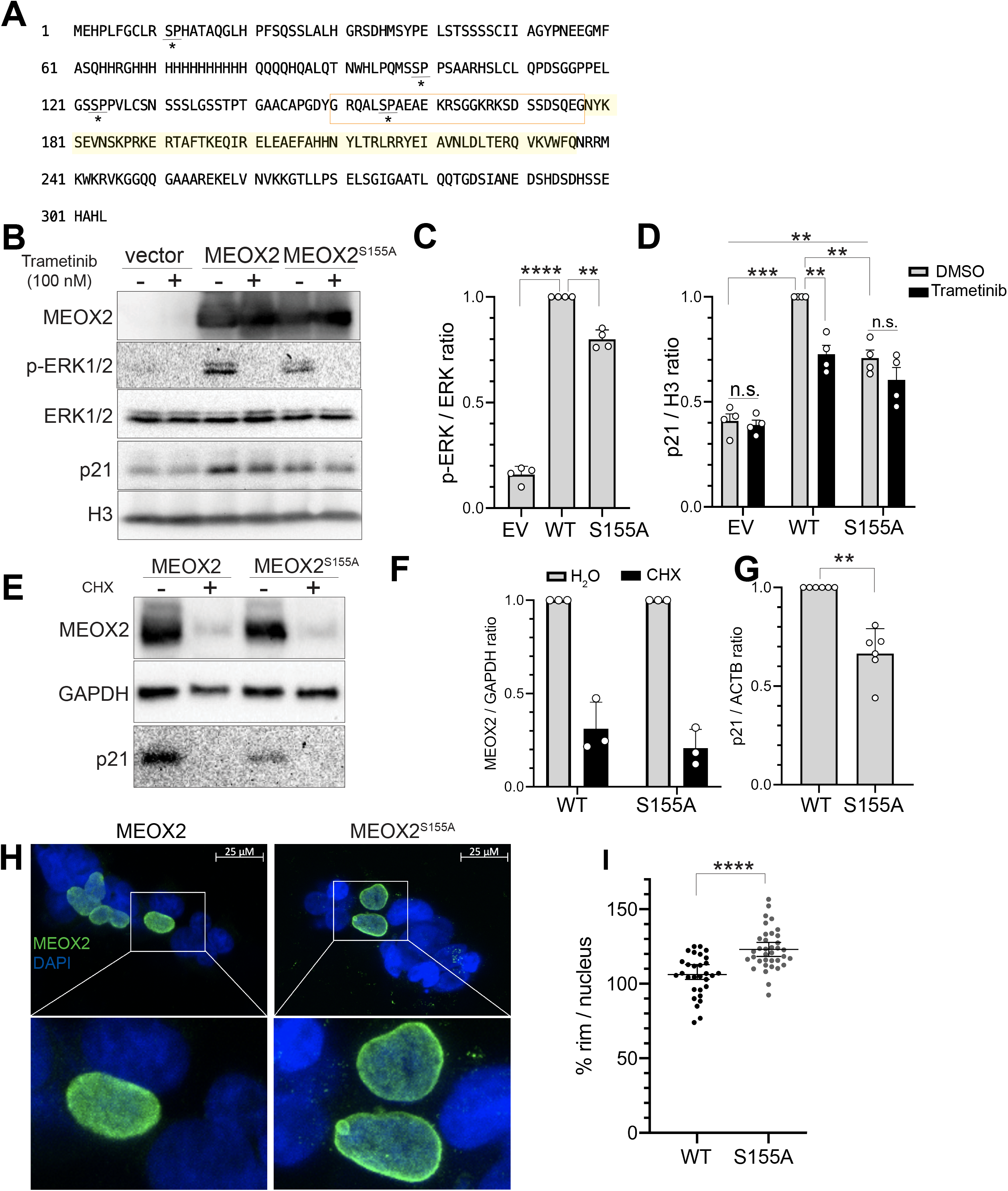
Phosphorylation of S155 affects the transcriptional activity of MEOX2. (A) The amino acid sequence of MEOX2. Red boxed regions include the phosphorylated S155 residue identified by immunoprecipitation followed by mass spectrometry in HEK293TN and TS667 cells. S-P motifs, the putative ERK phosphorylation site, are underlined and marked with an asterisk (*). The homeobox domain of MEOX2 is highlighted in yellow. (B) Western blot of HEK293TN cells 48 h after transient transfection with empty vector (vector), MEOX2, MEOX2^S155A^ and 24 hours after treatment with 100 nM trametinib (+) or DMSO as control (-) (C) Quantification of **(B)** showing p-ERK relative to total ERK levels. EV, empty vector; WT, MEOX2; S155A, MEOX2^S155A^. Mean ± SD, *n* = 4, two-sided Welsh’s test. (D) Quantification **(B)** showing p21 relative to H3 levels in trametinib treated and control cells. EV, empty vector; WT, MEOX2; S155A, MEOX2^S155A^. Mean + SD, *n* = 4, two-sided Welsh’s test. (E) Western blot showing protein stability of MEOX2 and MEOX2^S155A^ in HEK293TN cells measured 24 h after cycloheximide (CHX) treatment (200 μg/mL). (F) Quantification of **(E)** showing MEOX2 and MEOX2^S155A^ levels normalized to GAPDH. Mean ± ± SD, *n* = 3, two-sided Welsh’s test. (G) Quantification of p21 levels normalized to actin levels in MEOX2 and MEOX2^S155A^ transfected HEK293TN cells. Mean ± SD, n = 6, two-sided Welsh’s test. (H) Localization of MEOX2 and MEOX2^S155A^ in HEK293TN cells 48 h after transient transfection. Immunofluorescence using MEOX2 antibody (green) and DAPI (blue). Insets are zoomed images. Scale bar = 25 μm. (I) Quantification of MEOX2 localization in HEK293TN cells 48 h after transient transfection with MEOX2 (*n* = 30 cells) versus MEOX2^S155^ (*n* = 36 cells). Localized signal in nuclear membrane (rim) divided by nuclear signal (nucleus) is shown. Mean ± SD, two-sided Welsh’s test.

### MEOX2 promotes a growth phenotype

To determine whether MEOX2 expression regulates tumor growth *in vivo*, we identified two patient-derived GBM tumorsphere lines (L0125 and L0512) with low endogenous MEOX2 levels (Supplemental Fig. 3A). These lines were engineered to overexpress MEOX2, labeled with luciferase for *in vivo* bioluminescence (BLI) tracking, and implanted into the striatum of immunodeficient nude mice. BLI measurements showed that MEOX2 overexpression in L0125 induced a significantly faster growth phenotype than mice implanted with empty vector expressing cells, whereas no difference in growth kinetics was observed in the MEOX2 overexpressing L0512 line (Fig. 3A, B; Supplemental Fig. 3B-D).

**Figure 3.**
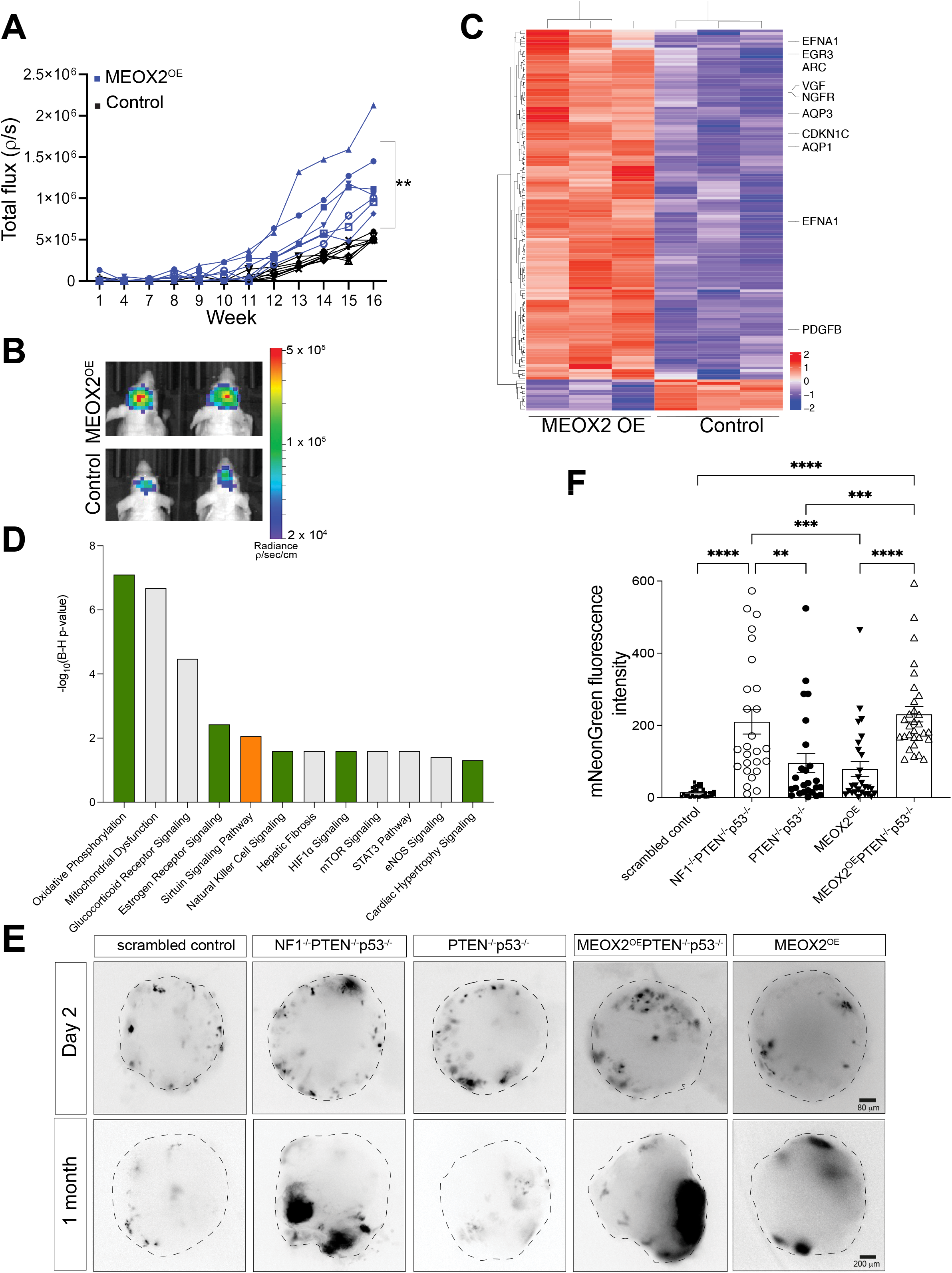
MEOX2 increases *in vivo* tumor growth and leads to increased proliferation in human cerebral organoid models of GBM. (A) Serial bioluminescence (BLI) imaging of the growth of L0125 GBM tumorsphere line transduced with control or MEOX2 (MEOX2^OE^) (7 mice in each group). BLI of each individual mouse is plotted. Mean ± SEM, two-sided Welsh’s test. (B) Example images of BLI in (A) are shown for mice implanted with MEOX2^OE^ (top) or control (bottom) cells. (C) Heatmap of differentially expressed genes (DEGs) in immortalized human astrocytes (IHAs) transiently transfected with MEOX2 compared to empty vector controls. (D) Enriched pathways in DEGs shown in **(C).** Colors of the bars indicate predicted activation states of the pathways identified by the Ingenuity Pathway Analysis (IPA). Green, activated; orange, inhibited; grey, unknown; B-H, Benjamini-Hochberg. (E) Images of cerebral organoids following nucleofection with scrambled control, NF1^-/-^PTEN^-/-^ p53^-/-^, PTEN^-/-^p53^-/-^, MEOX2^OE^PTEN^-/-^p53^-/-^or MEOX2^OE^ plasmids. Black color indicates mNeonGreen positive cells. Growth measured after 2 days (top row) and one month after nucleofection (1 month). OE, overexpression. (F) Quantification of mNeonGreen fluorescence intensity in cerebral organoids shown in (E). Control (*n* = 32), NF1^-/-^PTEN^-/-^p53^-/-^ (*n* = 26), PTEN^-/-^p53^-/-^ (*n* = 25), MEOX2^OE^ (*n* = 27), MEOX2^OE^PTEN^-/-^p53^-/-^ (*n* = 30). Only significant comparisons are shown (\**p*-value < 0.05). Oneway ANOVA, Tukey’s multiple comparisons test.

The heterogeneous *in vivo* growth response of patient-derived GBM lines to MEOX2 overexpression could be due to the faster basal growth kinetics of L0512 *in vivo*, suggesting that MEOX2 may increase tumor growth in GBM lines with slow basal *in vivo* growth dynamics such as L0125. Based on this hypothesis we wanted to test whether MEOX2 may be involved in tumor initiation and if this role may be mediated via target gene expression that could be usurped by constitutive RTK signaling in GBM. To determine which genes may be deregulated by MEOX2 in non-transformed cells, we transiently overexpressed MEOX2 in immortalized human astrocytes (IHA) and performed RNA-seq. We identified significant upregulation of 136 and downregulation of 12 genes (adjusted *p*-value < 0.05, absolute log_2_ fold-change >0.5) (Fig. 3C). Pathway analysis showed enrichment of several signaling networks, including STAT activation and HIF1α signaling (Fig. 3D). Amongst the significantly upregulated genes were *NGFR* as well as several members of the NGF-stimulated transcription (*VGF, EGR3, ARC*) and hypoxia (*EFNA1, CDKN1C, PDGFB, PKP1*) pathways (Fig. 3C; Supplemental Table 4). NGFR is highly expressed in GBM and inhibits the transcriptional activity of p53 to exert its oncogenic function (Zhou et al. 2016). Interestingly, *AQP1* and *AQP3*, members of the aquaporin (AQP) family, are upregulated upon MEOX2 overexpression. AQPs have been implicated in tumor cell growth and migration (Verkman et al. 2008).

We next aimed to determine whether MEOX2 might play a role in glioma initiation. To mimic tumor initiation *in vitro*, we used a cerebral organoid model to study whether overexpression of MEOX2 corroborates with the loss of canonical GBM tumor suppressors (PTEN, p53 and NF1). Organoids were electroporated with CRISPR-Cas9 to induce loss of indicated tumor suppressors together with stable overexpression of mNeonGreen with or without MEOX2 co-expression using a PiggyBac transposon system (Fig. 3E). This allows to monitor clonal outgrowth of genetically engineered fluorescently labeled cells in the context of a normal cerebral forebrain organoid. Remarkably, we found that MEOX2 synergized with p53 and PTEN loss to significantly increase clonal expansion of affected cells to the same level as the PTEN, p53 and NF1 triple knockout positive control (Fig. 3E, F). Compared to scrambled control, MEOX2 overexpression (adjusted p-value = 0.23) as well as p53 and PTEN loss alone (adjusted p-value = 0.08) did not lead to a significant increase in growth (Fig. 3E, F). These results suggest that MEOX2 in conjunction with additional mutations could act as an early driver of malignant transformation in gliomas.

### MEOX2 alters molecular pathways involved in tumorigenesis

To better understand the oncogenic role for MEOX2 at a molecular level, we performed transcriptomic analysis of two MEOX2 knockdown lines (TS600 and TS667) and two MEOX2 overexpression lines (L0125 and L0512) using RNA-seq. First, we analyzed the molecular changes in the MEOX2 KD lines compared to non-targeting controls (NTC) and identified the differentially expressed genes (adjusted *p*-value < 0.05, absolute log_2_ fold-change > 0.5) (Supplemental Table 5). In the TS667 line, 81 genes were up-regulated and 181 genes in both KD lines (Fig. 3A, top). In the TS600 line, MEOX2 KD2 led to a higher number of differentially expressed genes compared to KD1 (Fig. 3A, bottom), but overall fewer genes were altered compared to the TS667 MEOX2 KD lines. Notably, several genes including *ALK, EGFR, NOTCH3* and *NRCAM* were differentially expressed in both TS667 and TS600 lines (Fig. 4B, C). EGFR, a hyperactivated oncogene in GBM, and the associated signaling axis activate transcription factor networks that relay the oncogenic signals (An et al. 2018). This raises the intriguing possibility that MEOX2 could regulate EGFR expression in GBM. In addition, Notch signaling is deregulated in malignant gliomas (Bazzoni and Bentivegna 2019), and *NOTCH3* has been identified to drive cell motility and mesenchymal gene programs in neuroblastoma (van Nes et al. 2013). Furthermore, *VGF* is transcriptionally induced by MEOX2, as evidenced from IHA and TS667 RNA-seq data. VGF has been shown to trigger EMT and confer resistance to tyrosine kinase inhibitors (Hwang et al. 2017).

**Figure 4.**
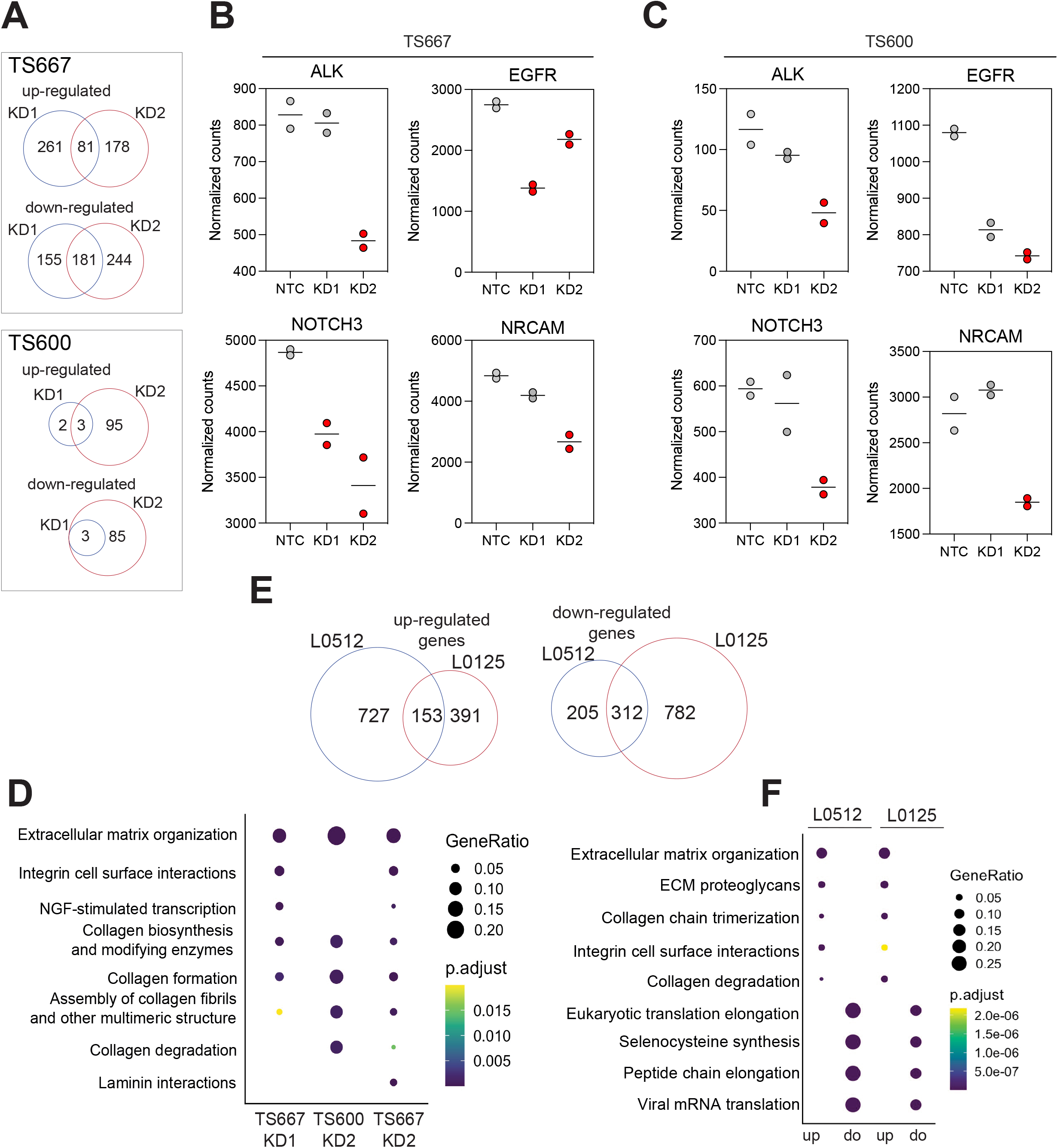
MEOX2 mediates activation of oncogenic pathways. (A) Venn diagrams showing overlap between up- or down-regulated differentially expressed genes (DEGs) in KD1 and KD2 TS667 (top) or TS600 (bottom) lines. KD, knockdown. (B) Normalized counts of ALK, EGFR, NOTCH3 and NRCAM in TS667 cells. Red dots indicate differential expression compared to NTC. NTC, non-targeting control. (C) Normalized counts of ALK, EGFR, NOTCH3 and NRCAM in TS600 cells. Red dots indicate differential expression compared to NTC. NTC, non-targeting control. (D) Significantly enriched pathways in TS667 KD1, TS600 KD2 and TS667 KD2 lines. Color indicates multiple-testing adjusted p-value of enrichment. (E) Venn diagram showing overlap between up-regulated (left) and down-regulated (right) DEGs in L0125 and L0512 MEOX2 overexpressing compared to empty vector expressing control lines. (F) Pathway enrichment among up- or down-regulated DEGs in L0125 and L0512 MEOX2 overexpressing compared to empty vector expressing control lines. Color indicates multiple-testing adjusted p-value of enrichment.

Next, we identified pathways and upstream regulators that were coherently altered upon MEOX2 loss. Several pathways, including hepatic fibrosis signaling, HOTAIR regulatory pathway, ILK signaling and GP6 signaling, and upstream regulators such as TGFB1 and AGT, were predicted to be less active (Supplemental Table 6). Several pathways including extracellular matrix organization and collagen biosynthesis were significantly enriched among downregulated genes upon MEOX2 loss (Fig. 4D).

Next, we identified the differentially expressed genes in MEOX2-overexpressing L0125 and L0512 GBM tumorsphere lines (adjusted *p*-value < 0.05, absolute log_2_ fold-change > 0.5) (Supplemental Table 7). In the MEOX2 overexpression lines, 153 genes were among the commonly upregulated and 312 downregulated genes in both lines (Fig. 4E). The upregulated genes were significantly enriched for several pathways, including extracellular matrix organization, integrin signaling pathway, and EMT (Fig. 4F). Taken together, these results suggest that MEOX2 may modulate ECM and collagens as part of its malignant gene expression program.

Our results indicate a context-dependent gene regulatory function of MEOX2 since although similar pathways are activated, not every cell line exhibits a similar molecular or phenotypic response to MEOX2 modifications. Such context-dependencies and paradoxical roles of transcription factors have been described in cancers(Blyth et al. 2005).

### Identification of direct MEOX2 target genes

As genome-wide binding patterns of MEOX2 are unknown, we performed antibody-guided chromatin tagmentation sequencing (ACT-seq) in TS600 and TS667 lines using KD2 as control. In addition, we performed cleavage under targets and tagmentation sequencing (CUT&Tag) in a primary fresh frozen GBM tumor. In total, 1519 peaks in TS667 and 1855 peaks in TS600 lines were identified compared to their respective KD2 and IgG controls (Fig. 5A; Supplemental Fig. 4A; Supplemental Table 8). Compared to IgG control, 2188 peaks were identified in the primary GBM tumor (Fig. 5B; Supplemental Table 8). In all samples, MEOX2 showed a distal binding occupancy, with most of the peaks located within intergenic and intronic regions (Fig. 5C). Peaks from both patient-derived tumorspheres and the GBM tumor harbored significant motif enrichment for putative MEOX1/2 sites (Fig. 5D; Supplemental Table 9). Interestingly, MEOX2 motif was highly ranked among the MEOX2 associated peaks in the TS667 and GBM compared to the TS600 line. Given our results that, in the TS600 line, p-ERK levels are unchanged upon MEOX2 loss and the overall number of differentially expressed genes are less compared to TS667 upon MEOX2 knockdown indicates that MEOX2 binding to its target genes may be weaker in the TS600 line.

**Figure 5.**
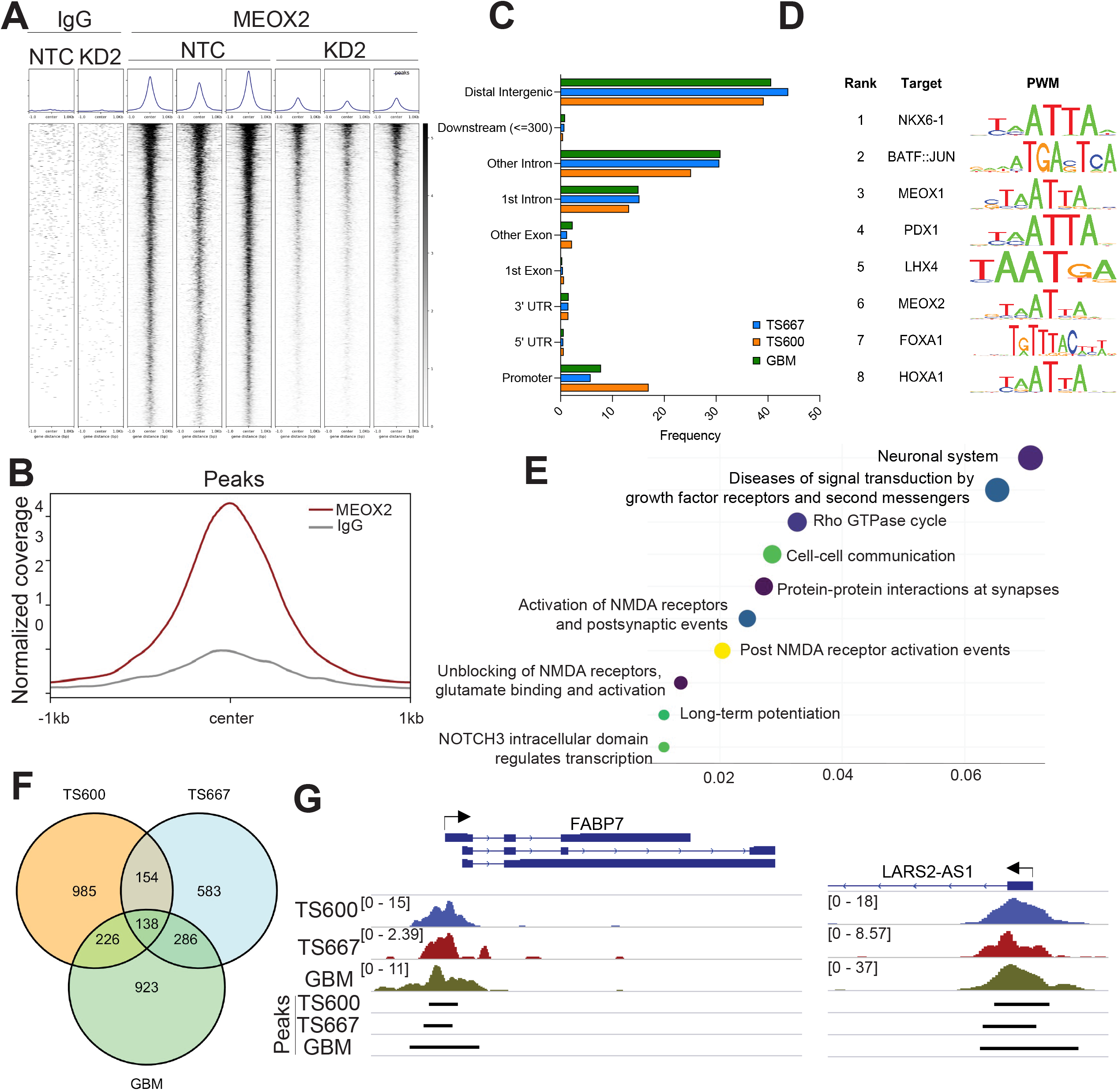
Identification and characterization of MEOX2 bound genomic regions. (A) Heatmap of normalized reads of genomic regions differentially bound by MEOX2 in TS600 line. MEOX2 peaks are ranked by intensity and shown relative to IgG negative controls in TS600 NTC and TS600 KD2 lines. Each experimental group includes 3 biological replicates. (B) Profiles of normalized reads of MEOX2-bound peaks in MEOX2 (dark red) and IgG (grey) CUT&Tag in a fresh frozen GBM sample. (C) Bar plot indicating the genomic feature distributions of the MEOX2-bound peaks in TS667 (blue), TS600 (orange), and GBM (green) samples. (D) Predicted sequence motifs and target genes for MEOX2-bound DNA sequences in TS600 line. PWMs, position weight matrices. (E) Pathway enrichment of MEOX2 peaks in the primary GBM tumor. Color indicates multiple-testing adjusted p-value of enrichment. (F) Venn diagram showing overlap of annotated genes identified from the differentially bound peaks in TS600, TS667 and primary GBM tumor. (G) Gene track view of normalized bigwig reads at the promoters of FABP7 (left) and LARS2-AS1 (right) of TS600 (blue), TS667 (dark red), and GBM (green) samples. The black lines indicate the called peaks corresponding to each sample.

To determine the biological functions of MEOX2 bound regions, we subjected the peaks to pathway enrichment analysis. This analysis identified several significantly enriched pathways, including PI3K/AKT signaling, MAPK signaling, cell-cell communication, and diseases of signal transduction (Fig. 5E; Supplemental Fig. 4B). A total of 138 genes were identified as bound by MEOX2 across all samples, including the ETS family (ETV1, ETV5, ETS1) and MAPK attenuators SPRY2 and DUSP10 (Fig. 5F). These results suggest that MEOX2 may potentially regulate the MAPK/ERK signaling by corralling and fine-tuning the ERK1/2 activity to prevent hyperactive ERK-associated toxicities to enable tumor initiation and growth in GBMs driven by RTK alterations. In addition, we overlapped the peaks in the tumorspheres and GBM tumor, which identified 44 overlapping peak-associated regions in all three samples (Supplemental Table 10). Among the overlapping genomic regions were several oncogenes such as *FABP7* and long non-coding RNAs such as *BHLHE40-AS1*, *LARS2-AS1* and *DLGAP1-AS1* (Fig. 5G). Notably, *FABP7* is downregulated upon MEOX2 loss, suggesting MEOX2 as a transcriptional activator of *FABP7* (Supplemental Table 5). Nuclear FABP7 is associated with infiltrative gliomas and poor prognosis in EGFR-overexpressing GBM (Liang et al. 2006).

Taken together, we identified MEOX2 as an oncogenic mesodermal transcription factor on chromosome 7 that activates several oncogenic pathways such as MAPK signaling that is central to GBM biology. We demonstrate that MEOX2 overexpression cooperates with loss of tumor suppressors (e.g. PTEN and p53) to drive significant growth in human cerebral organoid models of GBM, highlighting its role in tumor initiation. Given that MEOX2 is undetectable in the normal brain but co-opted and upregulated by GBM underscores the complexity of this disease along with the transcriptional networks hijacked by these tumors to maintain and amplify RTK signaling.

## Materials and Methods

### Patient sample

The fresh frozen GBM sample was obtained from a patient following surgical resection at the Department of Neurosurgery at the Acibadem Mehmet Ali Aydinlar University, Istanbul, Turkey. Use of patient material was approved by the Institutional Review Board at the Medical Faculty of Acibadem Mehmet Ali Aydinlar University. Informed consent was obtained from the patient included in the study.

### Cell Culture

Patient-derived GBM lines TS667, TS600, L0125, and L0512 were maintained in Neurocult Basal Medium with proliferation supplements, 20 ng/ml EGF, 20 ng/ml basic-FGF and 2 μg/ml Heparin (StemCell Technologies). L0125 and L0512 GBM tumorspheres were kindly provided by Dr. Rossella Galli (San Raffaele Scientific Institute, Milan, Italy). Immortalized human astrocytes were a gift of Dr. Russell O. Pieper (UCSF).

#### Cloning of CRISPR/Cas9 single guides

CRISPR/Cas9 single guides were designed using the online GPP sgRNA Design tool by Broad Institute in December 2017. The sequences of sgRNAs targeting MEOX2 are: sgRNA1 (KD1), gtcccgcgattatgcaagatg, sgRNA2 (KD2) gtgtgccgagccgcactcggt, and the sequence for the nontargeting control: accaatatcagtaata. LentiCRISPR v2 was digested with Esp3I at 37 °C for 4 hours, inactivated at 65 °C for 20 mins. For the digest, DTT was added to a concentration of 1mM. Digest with 6 x loading dye (NEB) was run on a 1 % agarose gel at 100 V for 1 h. Band was extracted and cleaned up using QIAquick Gel Extraction Kit (Qiagen). Guide RNA oligo pairs were annealed and phosphorylated using a reaction mix of 1μl of each oligo pair at 100μM, 1μL 10 x T4 ligation buffer, 0.5 μL T4 PNK added up to 10 μL reaction volume using water. Reaction was run in a thermal cycler at 37 °C for 30 mins, 95 °C C for 5 mins, ramped down to 25 °C at 0.1 °C/sec. Annealed oligos were diluted 1:500. Ligation was performed in a 1:5 ratio of digested and cleaned up back bone and annealed oligos. For reaction, 50 ng of vector were mixed with 1.5 μL of 10 x T4 ligation buffer, 1 μl T4 ligase and 2 μL diluted oligos (1:500) and filled up to 15 μL using water. Reaction was incubated at RT for 2 hours. 50 μL of Stbl3 chemically competent cells were transformed using 5 μL of the ligated reaction. Stabl3 were thawed on ice for 20 min, mixed with 5 μL of ligated reaction and incubated on ice for 20 min. Cells were then heat shocked using a heat block at 42 °C for 45 seconds and reaction tube was put back on ice for another 2 min. Stabl3 were grown for 45 min at 37 °C after addition of 250 μL SOC medium. Transformed Stbl3 cells were then plated on LB agar plates, containing 100 μg/mL ampicillin, and were kept at 37 °C overnight. Colonies were picked and plasmids were prepared using QIAprep Spin Miniprep Kit (QIAGEN). T7-endonuclease mismatch detection assay was performed following manufacturer’s protocol using Alt-R Genome Editing Detection Kit (IDT) to evaluate the activity of the site-specific nuclease.

#### Cloning of overexpression plasmids

MEOX2-FLAG or MEOX2 was cloned into pLVX-puro and MEOX2 was cloned into PB-linker-CAG_MCS_T2A_mNeonGreen for the cerebral organoid experiments. Empty plasmids were each double-digested for 4 h at 37 °C, using restriction enzymes XhoI and BamHI (NEB). Digests with 6 x loading dye (NEB) were run on a 1 % agarose gel at 100 V for 1 h. Bands were extracted and cleaned up using QIAquick Gel Extraction Kit (Qiagen). Inserts were amplified in a thermal cycler. Insert amplification was performed using the Kappa Hi Fi Hotstart ReadyMix (Roche). For a 25 μL reaction, 12.5 μL of Kappa Hi Fi Hotstart ReadyMix (Roche) and each 0.75 μL (0.3 μM) of 10 μM forward and reverse primers was pipetted into a reaction tube. 70 ng of template DNA were added, and reaction volume adjusted to 25 μL using PCR-grade water. Amplifications were followed by column clean ups using QIAquick PCR Purification kit (Qiagen). Amplified inserts were each double-digested for 4 h at 37 °C, using restriction enzymes XhoI and BamHI (NEB). Like backbones, inserts were run on gel and cleaned up using QIAquick Gel Extraction Kit (Qiagen). Ligations were performed using Quick Ligation Kit (NEB). Reaction mixes used were: 2 μL Quick Ligation Reaction Buffer (10 x), 25 ng (1:3 reaction) or 17 ng (1:5 reaction) of backbone DNA, 75 ng (1:3 reaction) or 83 ng (1:5 reaction) of insert DNA, 1 μL of Quick Ligase in a total reaction volume of 20 μL (adjusted with nuclease-free water). Reaction mixes were incubated for 5 min at RT. Following transformations of Stabl3 competent E-coli were performed using 5 μL of reaction mix per 50 μL competent bacteria.

### Viral transductions

TS667 and TS600 lines were infected with lentiCRISPRv2 encoding either nontargeting control or sgRNA targeting MEOX2 and selected with 0.5 μg/ml of puromycin. MEOX2 was stably overexpressed in L0125 and L0512 cell lines by viral transduction using pLVX-puro-MEOX2-FLAG or pLVX-puro-MEOX2-S155A-FLAG plasmid. Additional luciferase expression was stably induced by a second viral transduction using pLenti-PGK-V5-LUC Neo plasmid and selected with 400 μg/ml of G418. For virus production, HEK293TN cells were seeded in 10 cm dishes and transiently transfected with plasmid of choice using FuGene transfection reagent. Cells were triple-transfected with packaging and envelope plasmids in a 10:1 ratio, for virus production, and expression plasmid in a ratio of 1:1 to packaging plasmid. Cell culture medium was changed after 24 hours and collected for harvesting at 48, 56 and 72 hours after transfection. After the third harvest, the cell culture supernatant was mixed to a concentration of 10 % with PEG solution and incubated overnight at 4 °C. After 12 hours, the PEG/medium mix was centrifuged at 4000 x *g* for 45 min for virus collection. Supernatant was discarded and virus pellets were diluted in PBS. To select transduced cells, antibiotic was added in a concentration of 0.5 μg/mL for puromycin selection and 400 μg/mL Geneticin for neomycin selection every three days for 12 days.

### Site-directed mutagenesis

Ser^155^ of MEOX2 was mutated using using QuikChange Site-Directed Mutagenesis Kit (Agilent). The mutagenic primer was CCGCCAGGCACTGGCACCTGCGGAGGC, which converted Ser^155^ to alanine (S155A). The desired mutation was confirmed by Sanger sequencing.

### Cycloheximide stability assay

Cells were seeded and transiently transfected as stated before. For analysis of protein stability, cycloheximide (Selleckchem) was added in a concentration of 200 μg/mL to inhibit protein synthesis.

### Trametinib treatment

Cells were washed and seeded in DMEM-high glucose 10% FBS. Cells were transfected 24 h after seeding. Medium was changed to inhibitor containing medium 24 h after transfections. Cells were harvested after 24 h of inhibitor treatment.

### SDS page and Western Blot

Cells were directly lysed in the wells using 100μL–200μL of NP40 lysis buffer containing 1% Halt Protease and Phosphatase Inhibitor Cocktail (ThermoFisher). Lysates were collected and directly boiled at 95 C for 5 minutes for Western Blot (lysis buffer used) or used for BCA Protein Assay (NP40 lysis buffer used) according to manufacturer’s protocol (Thermo Fisher) followed by Western Blot. SDS gels were casted with a concentration of 10 % polyacrylamide. Equal amounts of protein, defined by BCA assay when using NP40 buffer, and defined by cell seeding numbers when using lysis buffer, were loaded on 10% polyacrylamide gels. Blotting was performed using the Mini-PROTEAN Tetra electrophoresis wet blot system (Bio-Rad) at 100 mV for 90 min or by using the iBlot 2 Dry Blotting System system (ThermoFisher) for 7 min transfer at 60 mV for mixed protein sizes. Membranes were blocked for 30 min in 5 % milk (v/v). Primary antibodies were applied overnight at 4 °C in 1% milk. Protein signal was detected using Pierce ECL Western Blotting Substrate (Thermo Fisher) and ChemiDoc XRS+ Gel Imaging System (Bio-Rad). If further probing on same membranes was performed, membranes were reconstituted using Restore PLUS Western Blot-Stripping-Buffer (ThermoFisher) for 2 times 15 minutes, followed by a TBST washing and incubation of membranes in 5 % milk before adding the next primary antibody ON at 4 °C.

For EGF stimulation experiments, TS667 and TS600 cells were washed and seeded in neural stem cell media without EGF/bFGF supplementation. Cells were kept for 48 h; afterwards medium was changed to neural stem cell media containing 50 ng/mL EGF. Cells were then collected at different timepoints (0 min, 10 min, 30 min, 2 h, 4 h, 6 h, 8 h) to observe pathway initiation by EGF treatment.

### Orthotopic transplantation

All mouse experiments were approved by the Institutional Animal Care and Use Committee at DKFZ. Female athymic Nude mice at age of 8 weeks (n=7 per group) were intracranially injected using a fixed stereotactic apparatus (Stoelting). Mice were weighed and injected with 5 mg/kg Carprofen subcutaneously. Mice were then put into an induction chamber with 1-3 Vol%. Once mice did not show reflex to toe-pinching, mice were put into stereotactic frame, on a heat mat on 36 °C, isoflurane was provided during following surgery via nose cone using 1-2 Vol% Isoflurane. Mouse eyes were covered with eye cream to prevent dryness. Mice were injected with 2 mg/kg bupivacaine near incision site for additional local anesthesia. Iodine solution was used on mouse head as a disinfectant for incision site. A small hole was drilled oriented 2.5 mm right of Bregma. Hamilton syringe was filled with 50,000 cells/μL cell solution. Injection of cells was performed using the previously drilled hole to access brain, syringe tip was positioned using stereotactic frame. 100,000 cells were injected 3 mm deep into the striatum at a speed of 1 μL/min. Cells were allowed to settle for 5 min before syringe was removed. Incision site was closed using tissue glue.

### Bioluminescence imaging

Cell lines for use in orthotopic *in vivo* experiments were labeled with pLenti-PGK-V5-LUC Neo (Addgene plasmid #21471). The 293 cells were seeded in 10-cm-diameter dishes and transfected with pCMV-dR8.2 dvpr (Addgene plasmid #8455), pCMV-VSV-G (Addgene plasmid #8454) and the reporter construct using Fugene (Promega, Madison, WI). Lentiviral particles were collected, filtered through a 0.45-μm syringe filter for infection. Transduced cells were selected using G418. Bioluminescent imaging was performed weekly following intraperitoneal injection of D-luciferin and measured using the Xenogen IVIS Spectrum *in vivo* imaging system (PerkinElmer). Living Image software (PerkinElmer) was used to acquire and analyze the BLI data.

### Cerebral organoid model

Human induced pluripotent stem cells (IPSCs) were treated with TripleLE dissociation reagent to obtain single cells, and were plated in a concentration of 9,000 cells/well into an ultra-low-binding 96-well plate in mTeSR™1 medium (Stemcell technologies) containing Rho-associated protein kinase (ROCK) inhibitor (1:200). The mTeSR™1 medium was changed every day and once the embryoid bodies (EB) reached a diameter size of around 500 μm, medium was replaced by cortical induction medium. After three days, EBs, now induced to COs, were pooled into a 6 cm dish containing fresh cortical induction medium. The next day, COs were nucleofected to induce tumorigenic events and amplification of MEOX2. In order to initiate the brain tumors in the COs, tumor suppressor knockouts of Tumor Protein 53 (TP53), Phosphatase And Tensin Homolog (PTEN) and neurofibromin (NF1) were introduced using CRISPR-mediated deletion. Each nucleofection reaction also contained a stably integrating piggybac plasmid, expressing mNeonGreen with or without additional MEOX2 expression. Briefly, 8-10 EBs were collected, resuspended in nucleofection reagent (Nucleofector™ kits for human stem cells P3, Lonza) containing plasmids and transferred into nucleofection vials. Nucleofection was performed according to the manufacturer’s protocol using the NucleofectorTM X-Unit (Lonza) with the electroporation pulse program CB-150. After nucleofection, COs were carefully transferred to one well of a 24-well plate containing cortical induction medium and cultured at 37 °C incubator overnight. Then, nucleofected COs were embedded into Geltrex and transferred to a low-attachment 6 cm dish containing cortical induction medium and incubated at 37 °C. One day after embedding, COs were transferred to the orbital shaker. From the following day, medium was replaced by differentiation medium with regular medium change every three to four days. Organoids were cultured for four weeks when mNeonGreen intensity was measured. After four weeks, organoids were imaged using a fluorescence microscope (Leica) and the Leica Las X software. For each organoid, three images in different Z-dimensions were acquired. The ImageJ software was used to measure the mNeonGreen intensity of each organoid by calculating the corrected total cell fluorescence (CTCF). After four weeks, organoids were imaged using a fluorescence microscope (Leica) and the Leica Las X software. For each organoid, three images in different Z-dimensions were acquired. The ImageJ software was used to measure the mNeonGreen intensity of each organoid by calculating the corrected total cell fluorescence (CTCF). This method determines the corrected total fluorescence by subtracting out background signal, which is useful for comparing the fluorescence intensity between different organoids. Briefly, all three images of one organoid were stacked and maximum intensity projection was performed. Within the stacked organoid image, all fluorescent areas were outlined and integrated density and mean grey value were measured. Afterwards, for background subtraction, three small areas within the same image were selected that have no fluorescence and the mean fluorescence of background readings was calculated. Finally, the CTCF value was calculated using the following formula: CTCF = Integrated Density – (Area of Selected Cell x Mean Fluorescence of Background readings). The CTCF value for each organoid was normalized to the whole area of the corresponding organoid.

### Immunofluorescence

Cells were seeded in a concentration of 5.000 cells/well in 24-well dishes in 500 μL of medium. Organoids were embedded in Tissue-Tek O.C.T. Compound (Sakura) and cut using a cryotome into sections of 20 μm and collected on SuperFrost (ThermoFisher) cover slides. For the staining, 24 h after seeding, medium was removed, and cells were washed twice using PBS. 400 μL 4 % PFA was added to each well and incubated for 20 min at room temperature. PFA was removed and cells were washed twice using PBS. Cells were permeabilized using 0.1 % of Triton X-100 for 15 min at room, washed three times using PBS, followed by three gentle washes using washing buffer. Cells were blocked using 400 μL of blocking buffer for 45 min at room temperature. Finally, primary antibody was diluted in dilution buffer and added to the cells and incubated at room temperature for 1 h. Primary antibody was removed after incubating for 1 h and cells were washed twice using wash buffer. Secondary antibody was added to the cells in 400 μL dilution buffer and incubated for 1 h at room temperature. Afterwards, cells were washed twice using wash buffer. Coverslips were mounted to microscopy slides using 1 drop of mounting media containing DAPI (VECTASHIELD) per coverslip.

### RNA-sequencing

TS667, TS600, L0125, L0512, IHA cells were seeded in a concentration of 5 x 10^5^ cells/well in 6-well dishes and harvested 48 h after seeding, during growth phase. IHA cells were seeded in a concentration of 5 x 10^5^ cells per well in 6-well dishes and transiently transfected using MEOX2-OE or control-plasmid. Cells were harvested 48 h after transfection. Total mRNA from biological triplicates of TS667-NTC, TS667-KD1, TS667-KD2, TS600-NTC, TS600-KD1, TS600-KD2 or L0125-MEOX2, L0125-control or L0512-MEOX2, L0512-control or IHA-MEOX2, IHA-control was extracted using the QIAGEN RNeasy RNA isolation kit according to manufacturer’s protocol. Extracted RNA was tested for quality using the Bioanalyzer system by Agilent, following manufacturer’s instructions. The sequencing of samples was performed by the DKFZ genomics and proteomics core facility. RNA quantity and quality were validated using 4200 TapeStation (Agilent), and samples with above an RNA integrity number of 9.8 were used for sequencing. RNA-seq libraries were created with TruSeq Stranded mRNA Library Prep Kit (Illumina). The sequencing performed was HiSeq 4000 Single-read 50 bp. The aligned bam files and feature counts were generated by the Omics IT and data management core facility at DKFZ (ODCF). Normalization of the raw counts and differential expression analysis was performed using *DESeq2* R package (Love et al. 2014). An absolute log_2_ fold-change of 0.5 and adjusted p-value of 0.05 was used to identify the differentially expressed genes in IHAs. Volcano plots were plotted using the R package *EnhancedVolcano* (Blighe et al. 2018). Enriched pathways were determined using the Ingenuity Pathway Analysis (IPA) software or the R packages *ReactomePA*(Yu and He 2016) and *clusterProfiler* (Yu et al. 2012).

### Antibody-guided Chromatin Tagmentation sequencing (ACT-seq) and Cleavage Under Targets and Tagmentation (CUT&Tag) sequencing

ACT-seq was largely performed according to Carter et al (Carter et al. 2019) as described in detail recently (Liu et al. 2021; PMID: 34070078) on triplicates of TS600 NTC and KD2 cells. For CUT&Tag experiments, nuclei isolation was performed according to the published protocol (Narayanan et al. 2020) using a minimum of 10^5^ nuclei. Isolated nuclei were washed and centrifuged at 600 *g* for 4 minutes. Nuclei were resuspended in 200 ul antibody buffer containing 4 ug MEOX2 antibody or IgG for control and incubated at 4 °C overnight. The Hyperactive In-Situ ChIP Library Prep Kit for Illumina (pG-Tn5) (Vazyme) was used for CUT&Tag. The next day, cells were centrifuged and washed once using Dig-wash buffer. Nuclei were then incubated in 200 μl Dig-wash buffer containing 2 μg of secondary antibody and incubate at RT for 30 min. Cells were washed three times using 200 μL of Dig-wash buffer. Finally, cells were resuspended in Dig-300 buffer and counted. For following steps, nuclei were diluted to a concentration of 100,000 cells/100 μl Dig-300 buffer and 0.8 μl pA-Tn5 were added to the mix and incubated at RT for 1 h. Followed by three wash steps using Dig-300 buffer. 40 μl tagmentation buffer were added to the mix and was incubated in a thermal cycler for 1 h at 37 °C. 10 μl of the diluted sample (≈10.000 nuclei), were mixed with 65 μl EB buffer, 0.5 μl 10% SDS and 0.5 μl Proteinase K. Mix was incubated at 50 °C for 1 h. Samples were purified using Ampure beads in a 1:1.4 ratio, incubated for 5 min at RT and placed on magnet followed by two wash steps using 200 μl of EtOH and 30 sec of incubation each. Supernatant was discarded and sample elute in 20 μl EB buffer. Next, bulk library was prepared using 30 μL of 2 x Kapa, 20 μL of bulk DNA, 5 μL of Index 1 (i5 10 μM) and 5 μL of index 2 (i7 10 μM) and run in a thermal cycler for 17 cycles. Concentrations were measured using Nano Drop. QC was performed using D1000 Tapestation.

### ACT-seq and CUT&Tag Analysis

The fastq files were trimmed using trimmomatic and aligned to hg19 with bowtie2 using parameters --end-to-end --very-sensitive --no-mixed --no-discordant. Mitochondrial and blacklisted regions were removed from the aligned bam files using samtools. Duplicates were removed using Picard tools (MarkDuplicates). Low quality reads (quality score < 2) were removed using samtools. Peaks were called using MACS version 2.1.2. Differentially bound sites were identified using *DiffBind* (R statistical package). For visualization, the bam files were normalized using BPM. The reads within peak regions were plotted as a heatmap using deepTools (Ramirez et al. 2014) and visualized using IGV tools. Specifically, coverage tracks were calculated with the following parameters: *--binSize 20, --normalizeUsing BPM, --extendReads 150*, using the tool bamCoverage. The peaks were visualized using the computeMatrix and plotHeatmap functions. Peaks were annotated using ChIPseeker R package.

### Immunoprecipitation and mass spectrometry

Whole cell lysates from three replicates per condition, each 5 x 106 cells, were prepared. 10 μg of MEOX2 antibody or 10 μg of IgG as a control were added to each of the samples. The reaction volumes were adjusted to 450 μL using NP40 lysis buffer. Reaction was incubated ON at 4 °C on a mixer. 25 μL of Pierce Protein A/G Magnetic Beads (Thermo Fisher) were placed into each of 12 (two conditions, three replicates, MEOX2 and IgG antibody samples) 1.5 mL Eppendorf tubes. 170 μL of Wash Buffer were added to the beads, tubes were rotated and after 5 minutes supernatants were removed while beads were collected on the sides of the tubes in a magnetic stand. Wash step was repeated using 1 mL of Wash Buffer in the second wash. Sample/antibody mixture from day 1 was added to the washed magnetic beads and incubated for 4 h at room temperature on a rotator. Beads were collected in a magnetic stand and flowthrough of lysate was saved for later analysis. Beads were washed five times each, using 500 μL of Wash Buffer per tube. Last wash step was performed using H2O instead of wash buffer. Lastly, protein of interest was removed from the Beads using 100 μL of Low pH Elution Buffer. Tubes were incubated for 10 minutes at room temperature and supernatant was collected in the magnetic stand and neutralized using 15 μL of Neutralization Buffer per tube. Proteomics were performed at the Mass Spectrometry Core Facility at the German Cancer Center (DKFZ).

### Antibodies

The following antibodies were used for all the experiments:

| Target | Source | Dilution | Company | Catalog no. | Method |
| --- | --- | --- | --- | --- | --- |
| **Primary antibodies** |  |  |  |  |  |
| MEOX2 | Rabbit | 1:1000,<br>10 $\mu\text{g}$ / 450 $\mu\text{l}$<br>(IP) | Atlas<br>Antibodies<br>(Sigma) | HPA053793 | Western<br>Blot,<br>IP, ACT-seq,<br>CUT&Tag,<br>IF |
| Phospho MAPK<br>(Erk1/2) | Rabbit | 1:2000 | Cell Signaling | #9101 | Western Blot |
| (Thr202/Tyr204) |  |  |  |  |  |
| MAPK (Erk1/2) | Mouse | 1:2000 | Cell Signaling | #9107 | Western Blot |
| P21 | Rabbit | 1:2000 | Cell Signaling | #2947 | Western Blot |
| GAPDH | Rabbit | 1:2000 | Cell Signaling | #2118 | Western Blot |
| Actin | Rabbit | 1:2000 | Cell Signaling | #4967 | Western Blot |
| H3 | Rabbit | 1:10000 | Cell Signaling | #9715 | Western Blot |
| **Secondary antibodies** |  |  |  |  |  |
| Rabbit IgG | Rabbit | 10 µg / 450 µL (IP) | Merck Millipore | PP64 | IP, ACT-seq |
| Alexa Fluor 488 | Donkey | 2 µg/mL | ThermoFisher | A-21206 | IF |
| Alexa Fluor 594 | Donkey | 2 µg/mL | ThermoFisher | A-21207 | IF |
| Anti-mouse | Horse | 1:5000 | Cell Signaling | #7076 | Western Blot |
| Anti-rabbit | Goat | 1:5000 | Cell Signaling | #7074 | Western Blot |

IP, immunoprecipitation; IF, immunofluorescence

### Statistical analysis

Two-sided t-test with Welsh’s correction was performed if not stated otherwise. Significance is indicated using following legend: ≥0.1234 (n.s.), ≤0. 0332 (*), ≤0.0021 (**), ≤0.0002 (***), ≤0.0001 (****).

### Accession Numbers

All data have been deposited in NCBI’s Gene Expression Omnibus under accession number GSE181146.

## Acknowledgements

We thank the members of the Turcan lab for helpful discussions. We thank the Genomics and Proteomics Core Facility (GPCF) at the DKFZ for providing next-generation sequencing (NGS) services and proteomics services and analysis. We thank the Omics IT and Data Management Core Facility (ODCF) at the DKFZ for data management and technical support. We thank the DKFZ Single-cell Open Lab (scOpenLab) for the support and experimental assistance. This work was supported by the German Cancer Aid, Max Eder Program grant number 70111964 (S.T.). J.T. is supported by a DFG Mercator Fellowship (DFG grant number TU 585/1-1). M.M and E.H. are supported by the Hector Stiftung II gGmbH.

## Author Contributions

A.S. and S.T. designed and directed the study. A.S., E.H., B.S., A.N. performed the experiments. A.S., E.H., D.W., M.M, S.T. analyzed the data. A.S., E.H., D.W., F.B, J.T., M.M., S.T. interpreted the data. J-W.P., M.B., C.S. provided technical assistance. D.K. provided a custom-built macro plugin for image analysis. E.H. performed the organoid experiments. M.M. supervised the organoid experiments. K.O. and M.N.P. provided the patient sample. M.B, D.W., C.P. provided expertise and supervised the ACT-seq experiments. F.B, W.W., C.P., J.T. and M.M. provided conceptual advice. A.S. and S.T. wrote the paper. All authors contributed to the writing and/or editing of the manuscript.

